# Unravelling the Physiological Correlates of Mental Workload Variations in Tracking and Collision Prediction Tasks: Implications for Air Traffic Controllers

**DOI:** 10.1101/2021.02.02.428702

**Authors:** Alka Rachel John, Avinash K Singh, Tien-Thong Nguyen Do, Ami Eidels, Eugene Nalivaiko, Alireza Mazloumi Gavgani, Scott Brown, Murray Bennett, Sara Lal, Ann M. Simpson, Sylvia M Gustin, Kay Double, Frederick Rohan Walker, Sabina Kleitman, John Morley, Chin-Teng Lin

**Affiliations:** Australian Artificial Intelligence Institute, Faculty of Engineering and Information Technology, University of Technology Sydney, Sydney, New South Wales, Australia; School of Psychology, University of Newcastle, Callaghan, New South Wales, Australia; School of Biomedical Sciences and Pharmacy and Centre for Advanced Training Systems, University of Newcastle, Callaghan, New South Wales, Australia; School of Life Sciences, University of Technology Sydney, Sydney, New South Wales, Australia; School of Psychology, University of New South Wales, Sydney, New South Wales, Australia; Centre for Pain IMPACT, Neuroscience Research Australia; Brain and Mind Centre and Discipline of Pharmacology, School of Medical Sciences, The University of Sydney, Sydney, New South Wales, Australia; Faculty of Science, The University of Sydney, Sydney, New South Wales, Australia; School of Medicine, Western Sydney University, Sydney, New South Wales, Australia

**Author notes:** **Corresponding Author:** Alka Rachel John. **Author Note** We have no known conflict of interest to disclose. Correspondence concerning this article should be addressed to Alka Rachel John, Australian Artificial Intelligence Institute, Faculty of Engineering and IT, University of Technology Sydney, NSW 2007, Australia.

**Keywords:** Mental workload, EEG, pupil size, blink rate, RMSSD

## Abstract

**Objective:** We have designed tracking and collision prediction tasks to elucidate the differences in the physiological response to the workload variations in basic ATC tasks to untangle the impact of workload variations experienced by operators working in a complex ATC environment.

**Background:** Even though several factors influence the complexity of ATC tasks, keeping track of the aircraft and preventing collision are the most crucial.

**Methods:** Physiological measures, such as electroencephalogram (EEG), eye activity, and heart rate variability (HRV) data, were recorded from 24 participants performing tracking and collision prediction tasks with three levels of difficulty.

**Results:** The neurometrics of workload variations in the tracking and collision prediction tasks were markedly distinct, indicating that neurometrics can provide insights on the type of mental workload. The pupil size, number of blinks and HRV metric, root mean square of successive difference (RMSSD), varied significantly with the mental workload in both these tasks in a similar manner.

**Conclusion:** Our findings indicate that variations in task load are sensitively reflected in physiological signals, such as EEG, eye activity and HRV, in these basic ATC-related tasks.

**Application:** These findings have applicability to the design of future mental workload adaptive systems that integrate neurometrics in deciding not just ‘when’ but also ‘what’ to adapt. Our study provides compelling evidence in the viability of developing intelligent closed-loop mental workload adaptive systems that ensure efficiency and safety in ATC and beyond.

**Précis:** This article identifies the physiological correlates of mental workload variation in basic ATC tasks. The findings assert that neurometrics can provide more information on the task that contributes to the workload, which can aid in the design of intelligent mental workload adaptive system.

## Introduction

People tend to avoid performing tasks that push their capabilities beyond their limits as they find it frustrating and stressful (Ahlstrom, 2010). However, not all work environments offer that luxury, which makes it crucial to establish good interaction between the human operator abilities and work environment (Wickens et al., 2015). Even though human operators can easily adapt to diverse work environments and perform several tasks and use different equipment simultaneously, poorly designed work environments cause an overload of sensory information resulting in excess workload. Air traffic controllers operate in such a complex environment to ensure a safe and efficient air traffic flow by organising traffic flow in a way that aircraft reach their destination in a well-organized and expeditious manner. However, as the air traffic increases, there is a growing need to study the mental factors that ensure the efficiency of air traffic controllers.

Mental workload is one of the most crucial factors that affects the efficiency of air traffic controllers as they operate in complex interactive work environments. Electroencephalogram (EEG) signal has been widely employed to estimate mental workload as the effects of task demand are clearly visible in EEG rhythm variations (Brookings et al., 1996, Gevins and Smith, 2003, Radüntz and Meffert, 2019). However, EEG features of the mental workload are found to be task-dependent, therefore, adding other modalities like eye activity data and heart rate data can help achieve far superior outcomes (Ke et al., 2014, Popovic et al., 2015).

Once the mental workload of the operator can be reliably assessed, it can be used to drive a mental workload adaptive system (Prinzel et al., 2000; Schmorrowe et al., 2006). A mental workload adaptive automation system should be able to conform to the variations in the mental workload of the operator without them having to explicitly state their needs or triggering the automation. When human operators and automation team up to achieve better performance and efficiency, the operator expects automation to behave like a human coworker (Aricò et al., 2017). Therefore, adaptive automation should be timely, stepping in at the right time and cognitively empathetic with the operator, helping where it is needed, taking over the task that is currently overwhelming the operator. However, currently, physiological correlates of the mental workload are only used to decide “when” to adapt and not “what” to adapt, keeping the strategies employed by the adaptive automation system still primitive. There is a need to develop intelligent adaptive systems that can identify what form of automation to use depending on the type of mental workload experienced by the operator. Nonetheless, there is still a dearth in evidence that physiological metrics of mental workload can direct to the tasks contributing to workload.

In this paper we investigated whether the multimodal physiological metrics of mental workload can provide more information about the task contributing to the workload experienced by the ATC operator. Even though several factors influence the complexity of ATC tasks (Mogford et al., 1995, Cummings and Tsonis, 2005), such as environmental, display, traffic and organisational factors, the main functions for ATC operator are tracking and collision prediction. Therefore, we designed tracking and collision prediction tasks to elucidate the physiological effects of workload variations in these basic ATC tasks. We formulated the following four research hypotheses for our study:

H1. The three distinct levels of workload defined in both tracking and collision prediction tasks can yield significant performance degradation with the increasing levels of workload.

H2. Workload variation in tracking and collision prediction tasks can be reliably assessed using EEG, eye activity and HRV metrics.

H3. The performance in tracking and collision prediction tasks can be predicted based on the measured physiological signals.

H4. Physiological response to the workload variations in the tracking and collision prediction tasks will be distinct across tasks.

## Methods

### Participants

Twenty-four participants (age 25 ± 5, 17 males and 7 females, all right-handed) participated in this experiment after giving written informed consent. The experimental protocol was approved by the University of Technology Sydney Human Research Ethics Expedited Review Committee (ETH19-4197).

The EEG data were collected using SynAmps2 Express system (Compumedics Ltd., VIC, Australia) with 64 Ag/AgCl sensors system. Eye activity data was collected using Pupil Labs Pupil Core (Berlin, Germany). The Blood Volume Pulse (BVP) data was recorded using Empatica E4 (Empatica Srl, Milano, Italy). The real-time synchronisation of events from the task scenario to the EEG, eye activity and BVP data was achieved by the Lab Streaming Layer (Kothe, 2015).

### Experimental Procedures

Our experimental design included two tasks – multiple objects tracking task (Innes et al., 2019) and collision prediction task. As shown in Figure 1(A), in the tracking task, during the initial 3 seconds, participants look at a fixation cross on the screen followed by a freeze phase, where the dots, some of which are blue, and the rest are red, remain stationary. The blue dots are the dots that need to be tracked (hence, ‘targets’). After three seconds of freeze, the blue targets also turn red so that they are no longer distinctive from the other dots and all the dots start moving. The participant is asked to keep track of the targets (dots that were initially blue) for 15 seconds. After this time window all dots stop moving and the participants should indicate the target dots by clicking on the dots that they have kept track of. The workload levels in this tracking task are manipulated by varying the number of blue dots and the total number of dots (see Table 1).

**Table 1:**
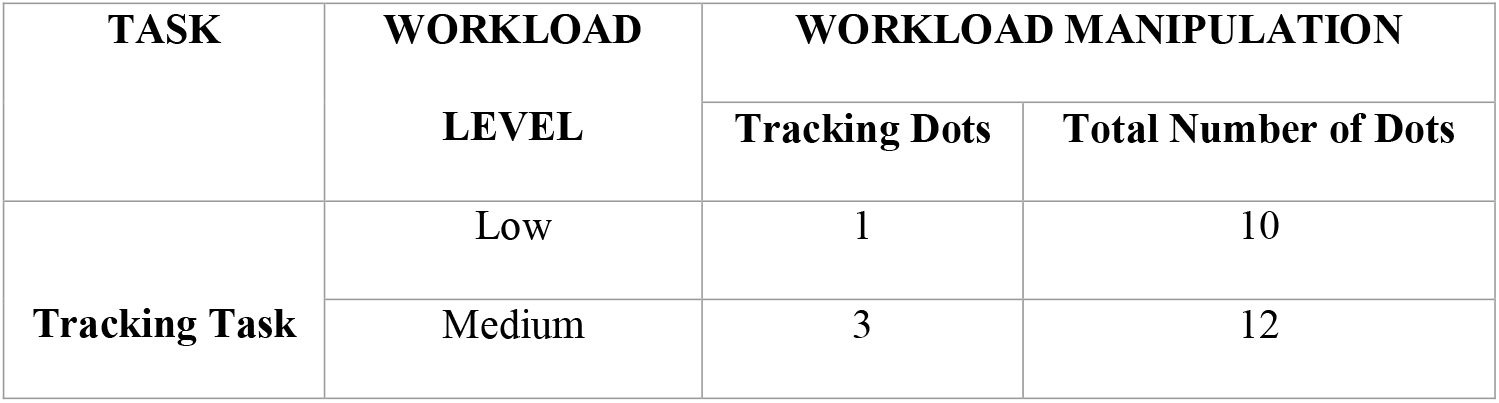
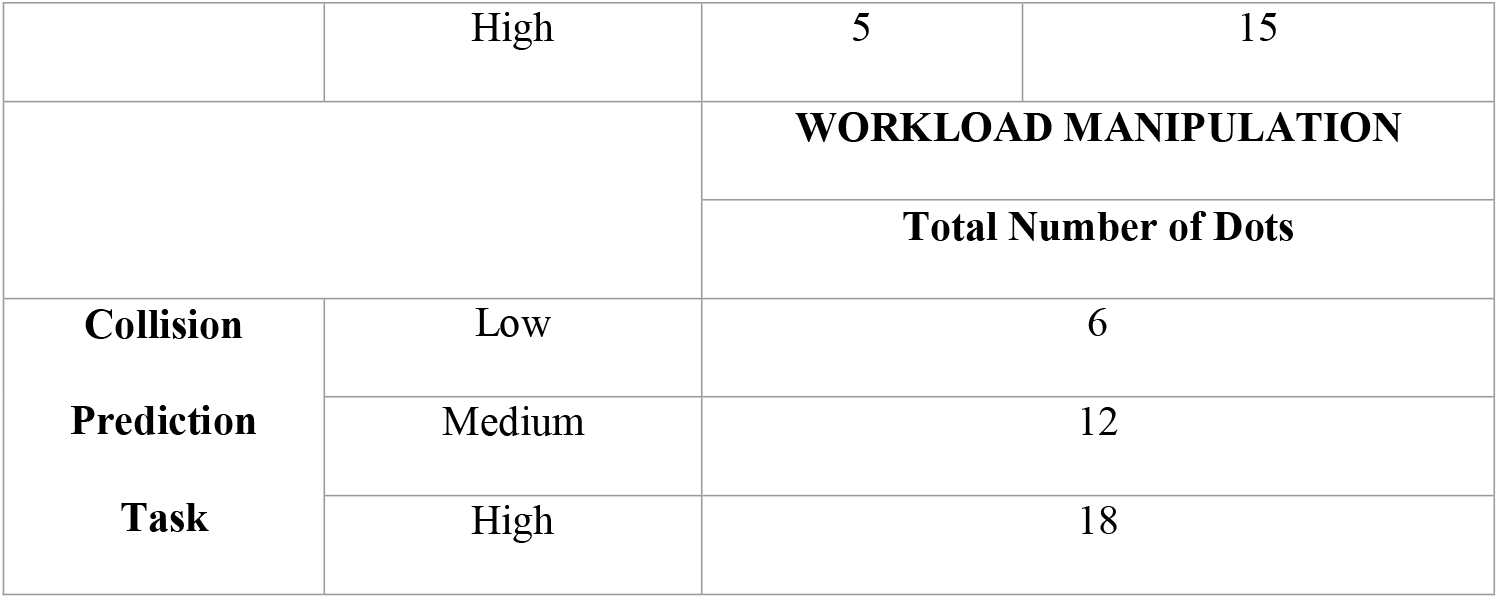
Workload Manipulations in the tracking and collision prediction tasks

**Figure 1:**
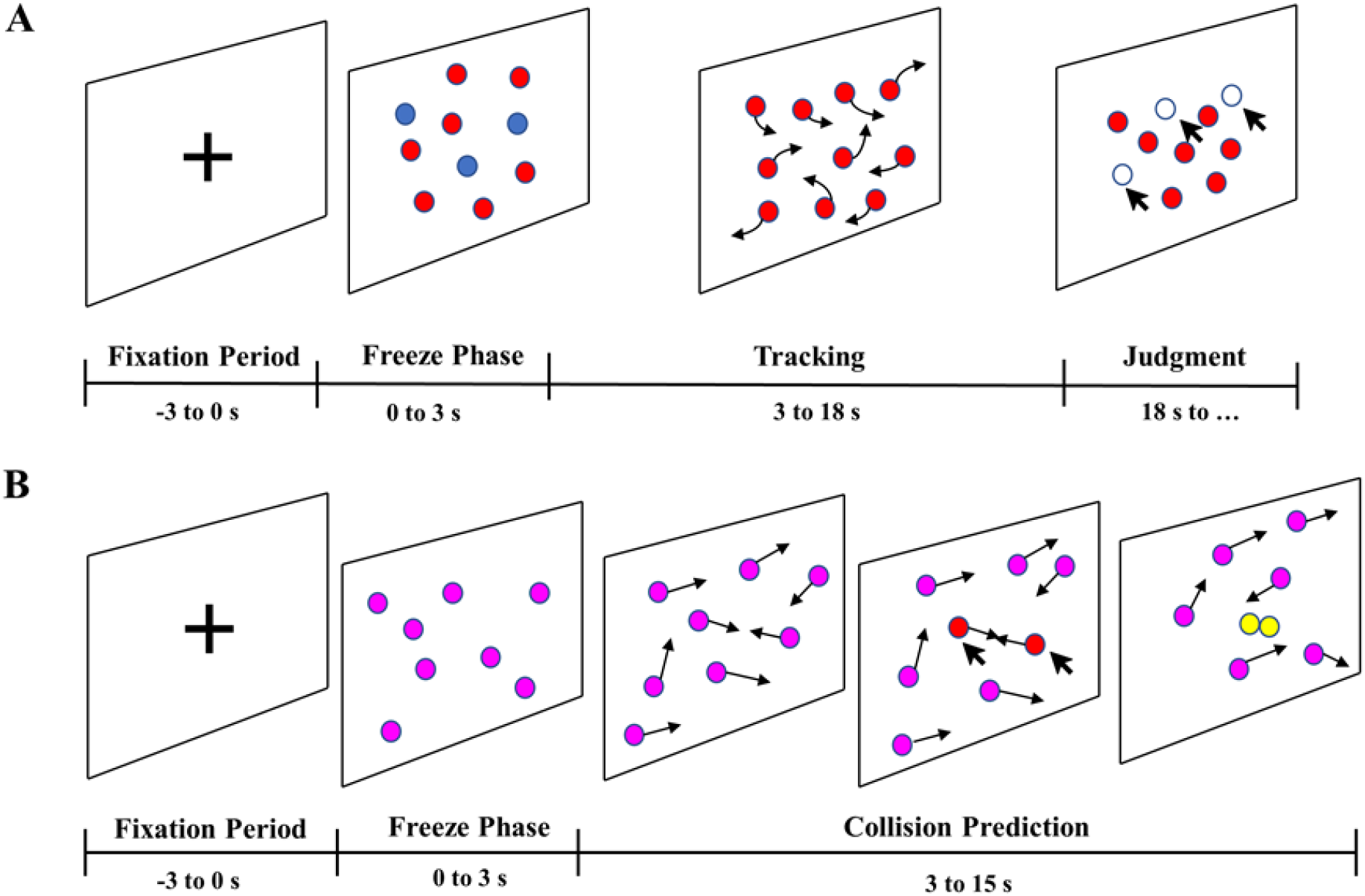
The experimental design of the tasks. (A) the experimental design of the tracking task and (B) the desig of the collision prediction task. The number of dots shown in these diagrams is just for representation purposes.

As shown in Figure 1(B), in the collision prediction task, there is a fixation cross on the screen for three seconds. Then there is a three-second-long freeze phase where the dots remain stationary, after which all the dots start moving. The participant is required to predict the trajectory of the dots and identify which pair of dots would collide. We have manipulated the trajectory of the dots such that there will be only one collision in each trial. The participants were asked to identify the pair of dots that would collide and click on both dots before the collision happens. The levels of workload were manipulated by varying the number of dots (see Table 1).

Each participant had to perform 108 trials of each task with 36 trials of each workload level. The type of workload condition in the trials was randomised to avoid any habituation or expectation effects. All participants were trained in a training session to familiarise themselves with th tasks. After the training, all participants performed the tasks for ∼ 1.5 hours during which EEG, eye activity and HRV data were collected.

### Data Analysis

#### Behavioural and Performance Data Analysis

For the tracking task, each participant’s performance was evaluated by examining the tracking accuracy.

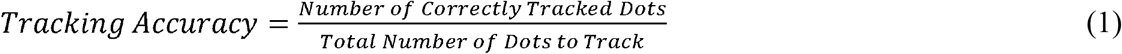

In case of the collision prediction trials, the performance was determined using the time before collision and collision miss proportion rate. The time before collision is the time period between when the participant clicks on either one of the colliding dots and when the collision happens (see Supplementary Figure 1). A collision miss was considered to happen when the participant was unable to identify which pair of dots would collide and, hence, did not click on either of the dots before the collision. The collision miss proportion rate for a particular workload level of the collision prediction task is the ratio of the number of collision prediction misses to the total number of collisions in that specific workload level.

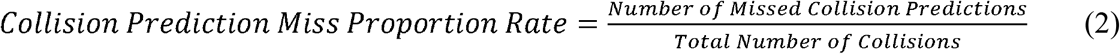

#### EEG Preprocessing

EEG data were preprocessed (see Supplementary Figure 2) using EEGLAB v2020.0 toolbox (Delorme and Makeig, 2004) in MATLAB R2019a (The Mathworks, Inc., Natick, MA, USA). EEG data were down-sampled to 250 Hz, and a band-pass filter of 2–45 Hz was applied. Channels with three seconds or more flat line were removed using the clean_flatline function. Noisy channels were identified and removed using the clean_channels function in EEGLAB. On an average 3±1 channels were removed and these channels were restored by interpolating the data from neighbouring channels using the spherical spline method from the EEGLAB toolbox. Continuous artifactual regions were removed using the EEGLAB function, pop_rejcont. Then window cleaning was performed using the clean_windows function in EEGLAB. After these artifact removal steps, two EEG datasets were extracted, one comprising tracking trials and one with the collision prediction trials. Each participant had 34±2 high workload, 35±1 medium workload and 34±1 low workload tracking trials, and 32±2 high workload, 33±2 medium workload and 33±1 low workload collision prediction trials.

**Figure 2:**
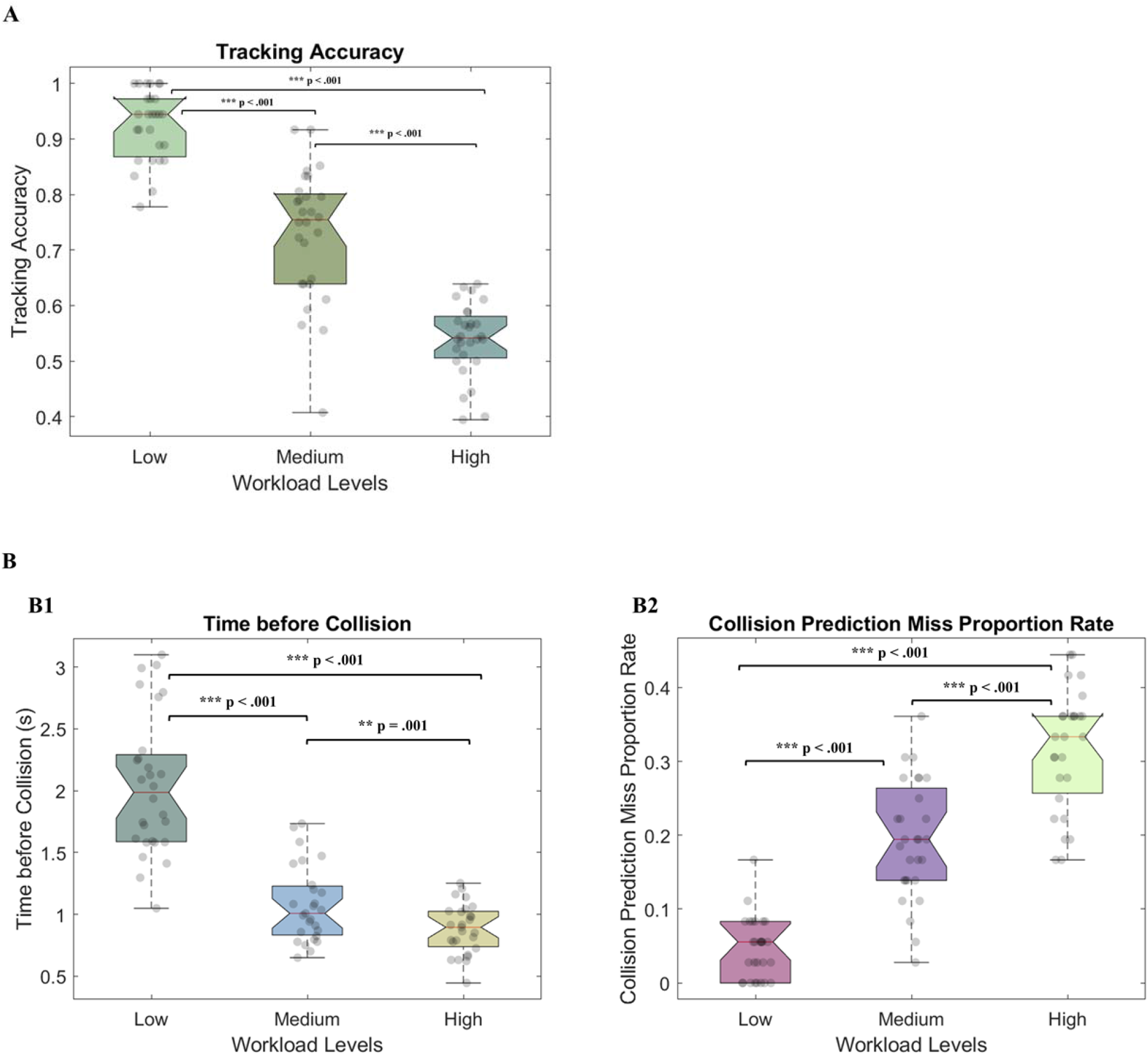
(A) shows the tracking accuracy of all the participants in the tracking task for the three levels of workload. (B) shows the performance of all participants in the collision prediction task for the three levels of workload. (B1) shows the mean time before collision for all the participants in the low, medium, and high workload conditions. (B2) shows the collision prediction miss proportion rate for the three levels of workload.

Tracking and collision prediction datasets were decomposed using Independent Component Analysis (ICA), performed using EEGLAB’s runica algorithm (Delorme and Makeig, 2004). Finally, we employed ICLabel (Pion-Tonachini et al., 2019), an automatic IC classifier to identify and reject components related to heart, line noise, eye, muscle, channel noise and other activities.

#### IC Clustering

EEGLAB STUDY structure (Delorme et al., 2011) was used to manage and process data recorded from multiple participants. A Study was created for each task, and each Study had one group (with 24 participants) with three conditions corresponding to the three levels of workload. For each participant, only those ICs that had a residual variance (RV) less than 15% and inside the brain volume were chosen, which was achieved using Fieldtrip extension (Oostenveld et al., 2011). The k-means clustering algorithm (Hartigan and Wong, 1979) was used to cluster independent components across all participants to clusters based on two equally weighted (weight = 1) criteria: (1) scalp maps and (2) their equivalent dipole model locations, which was performed using DIPFIT routines (Oostenveld and Oostendorp, 2004) in EEGLAB. Frontal and parietal brain regions have been reported to reflect the changes in workload (Brookings et al., 1996; Aricò et al., 2017), and as both our tasks also manipulate the visual load, we particularly focused on the frontal, parietal and occipital clusters of brain activity. Talairach coordinates (Lancaster et al., 2000) of the fitted dipole sources of these clusters were identified to select frontal, parietal and occipital clusters.

The grand-mean IC event-related spectral power changes (ERSPs) for each condition was subsequently calculated for each cluster. The three seconds of fixation phase in each tracking and collision prediction epoch was taken as the baseline to see the changes in power spectra during the task. ERSPs for frontal, parietal and occipital clusters for both tracking and prediction tasks were examined. To compare the ERSP of different workload conditions, permutation-based statistics, implemented in EEGLAB, was used with Bonferroni correction and significance level set to p = .05. Also, for the frontal, parietal and occipital cluster, each ICs’ spectral powers were calculated using EEGLAB’s spectopo function, which uses Welch’s periodogram method (Welch, 1967) on each 2-s segment using a Hamming window with 25% overlap for a range of frequencies from 2 to 45 Hz. For each IC, the power spectral density (PSD) at different frequency bands were examined to identify the correlates of mental workload.

#### Eye Activity data

Pupil Core software, Pupil Capture provides the pupil size for the left and right eye separately along with the associated confidence value, which represents the quality of the detection result. All data points where the confidence of the pupil size was less than 0.8 were removed from the data. The pupil size data was normalised using the baseline data (defined as the three seconds of fixation period in each tracking and collision prediction epoch). The number of blinks during each trial was also extracted from the pupil size measurement when the pupil size and confidence of the measurement, reported by the Pupil Capture software, suddenly dropped to zero.

#### Heart Rate Variability

Inter-beat-interval (IBI) time series was computed from the Blood Volume Pulse (BVP) data of each tracking and collision prediction trial. Root Mean Square of the Successive Differences (RMSSD) was computed by detecting peaks of the BVP and calculating the lengths of the intervals between adjacent beats.

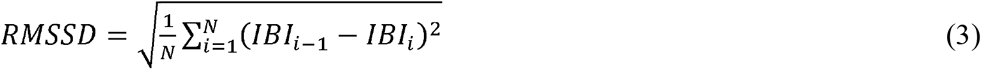

RMSSD data was also normalised by considering the three seconds of fixation period in each tracking and collision prediction epoch as the baseline.

### Statistical Analysis

Statistical analyses were carried out using the SPSS (IBM SPSS 26.0; Chicago, IL, U.S.A.) statistical tool. In order to investigate the differences in the performance, EEG, eye activity and HRV parameters across participants in the three workload levels of tracking and collision prediction tasks, one-way repeated-measures analysis of variance (ANOVA) was conducted with workload level as the within-subjects factor. Mauchly’s test was implemented to test for sphericity. We performed Greenhouse-Geisser correction if sphericity was not satisfied (p < .05). If the main effect of the ANOVA was significant, post-hoc comparisons were made to determine the significance of pairwise comparisons, using Bonferroni correction. Finally, multiple linear regression was performed to relate EEG, eye activity and HRV metrics to the performance in the tracking and collision prediction tasks. EEG power, eye activity and HRV metrics were all entered as predictors using the enter method, and the performance in the task was the dependent variable.

## Results

### Behavioural and Performance Measures

A repeated-measures ANOVA showed that tracking accuracy decreased significantly with increasing levels of workload [F(2, 54) = 239.910, p < .001, η_p_^2^ = .899], as shown in Figure 2(A).

For the collision prediction task, the time before collision and collision prediction miss proportion rate was considered. A repeated-measures ANOVA results showed that time before collision decreased significantly with increasing workload [F(1.497, 40.406) = 132.688, p < .001, η_p_^2^ = .831] and the collision prediction miss proportion increased with increasing levels of workload [F(1.593, 43.009) = 116.338, p < .001, η_p_^2^ = .812], as shown in Figure 2(B1) and 2(B2).

### EEG Results

#### Independent Source Clusters

The frontal, parietal and occipital clusters for both tracking (refer Figure 3) and collision prediction task (see Figure 4) were selected based on the location of fitted dipole sources (Oostenveld and Oostendorp, 2004).

**Figure 3.**
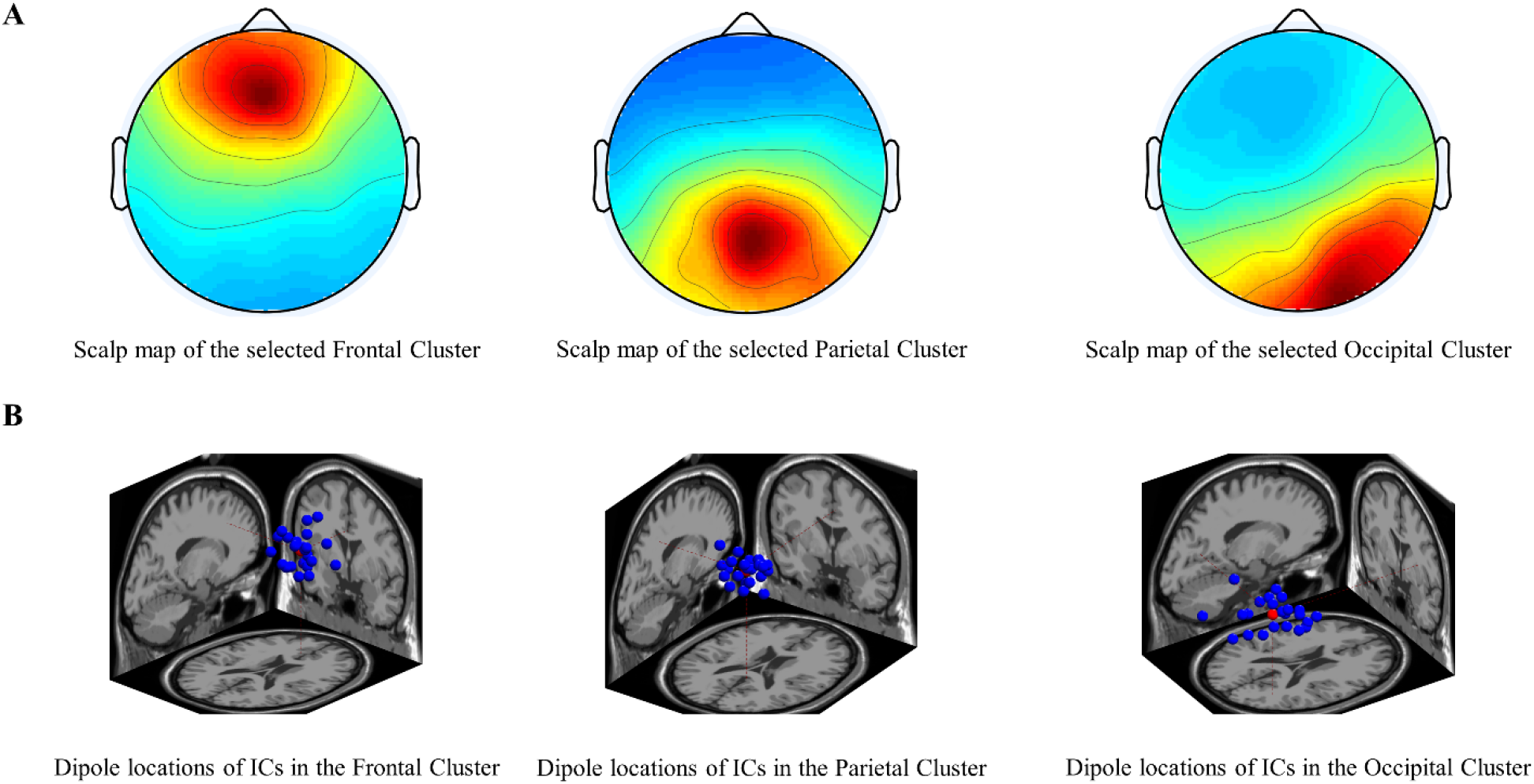
Frontal [Talairach coordinate: (−1, 41, 27)], Parietal [Talairach coordinate: (4, -51, 39)] and Occipital [Talairach coordinate: (30, -70, 15)] clusters selected in the tracking task (A) spatial scalp maps; (B) dipole source locations.

**Figure 4.**
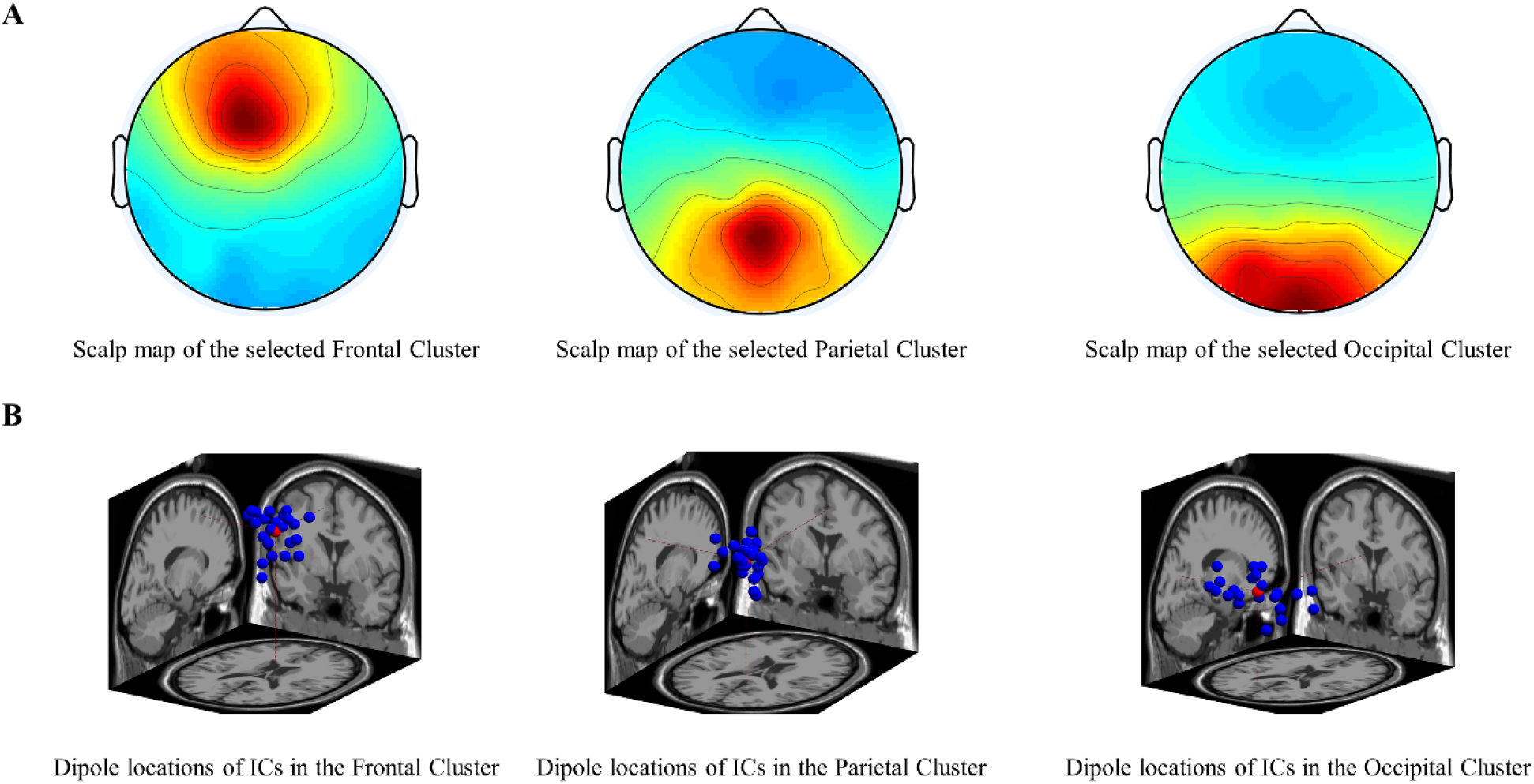
Frontal [Talairach coordinate: (−10, 17, 46)], Parietal [Talairach coordinate: (5, -47, 47)] and Occipital [Talairach Coordinate: (−3, -69, 20)] clusters selected in the collision prediction task (A) spatial scalp maps; (B) dipole source locations.

#### ERSP Changes with Mental Workload

Figures 5 illustrates frontal, parietal and occipital clusters’ ERSP changes for three workload conditions: low, medium and high during the tracking task. Statistical analysis on ERSP changes of the frontal cluster (Figure 5(A)) revealed a significant increase in theta power with increasing levels of workload. However, no significant spectral power variations were observed at the parietal cluster. Figure 5(B) shows the ERSP changes at the occipital cluster, which revealed a significant decrease in alpha power with increasing levels of workload.

**Figure 5:**
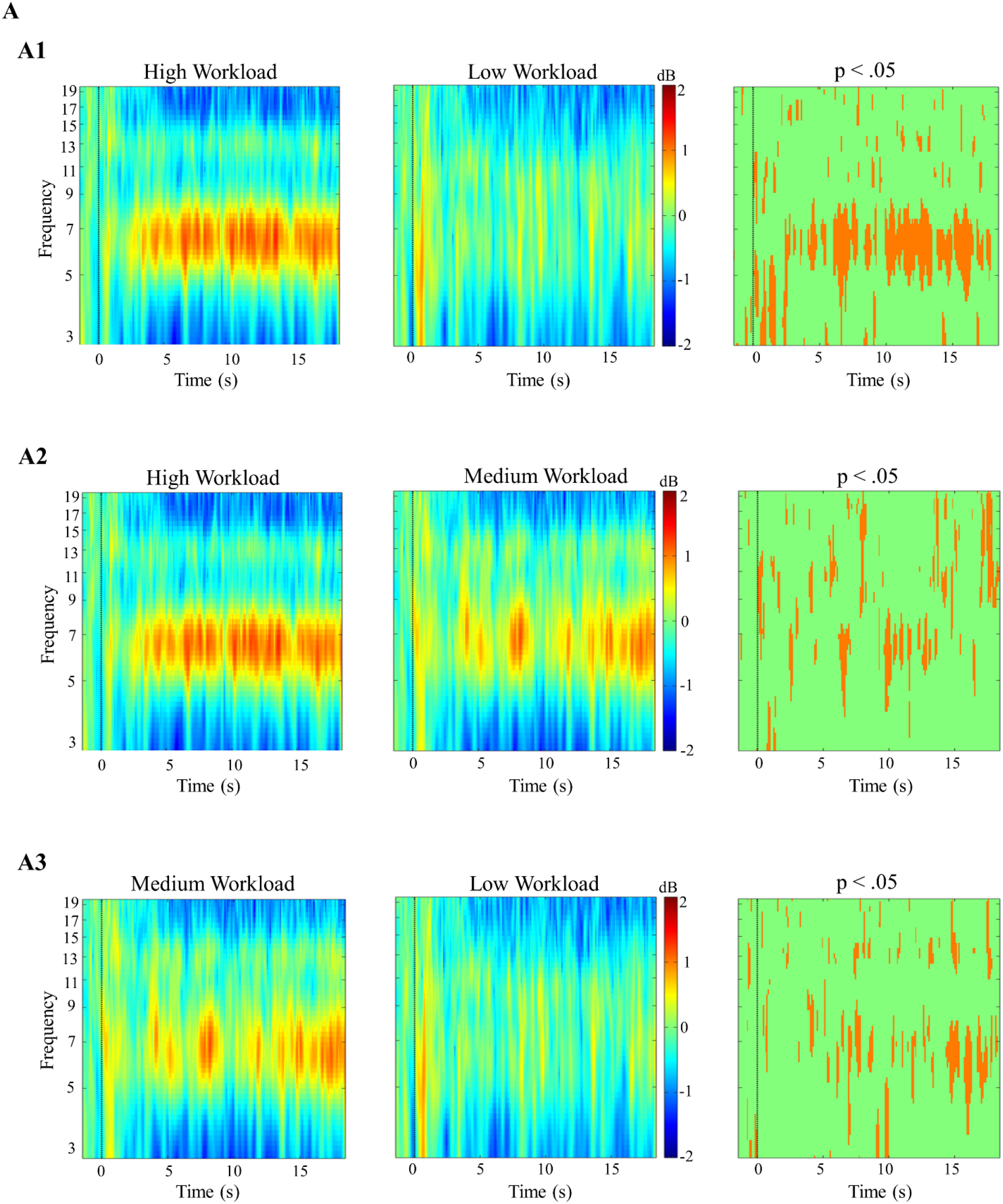
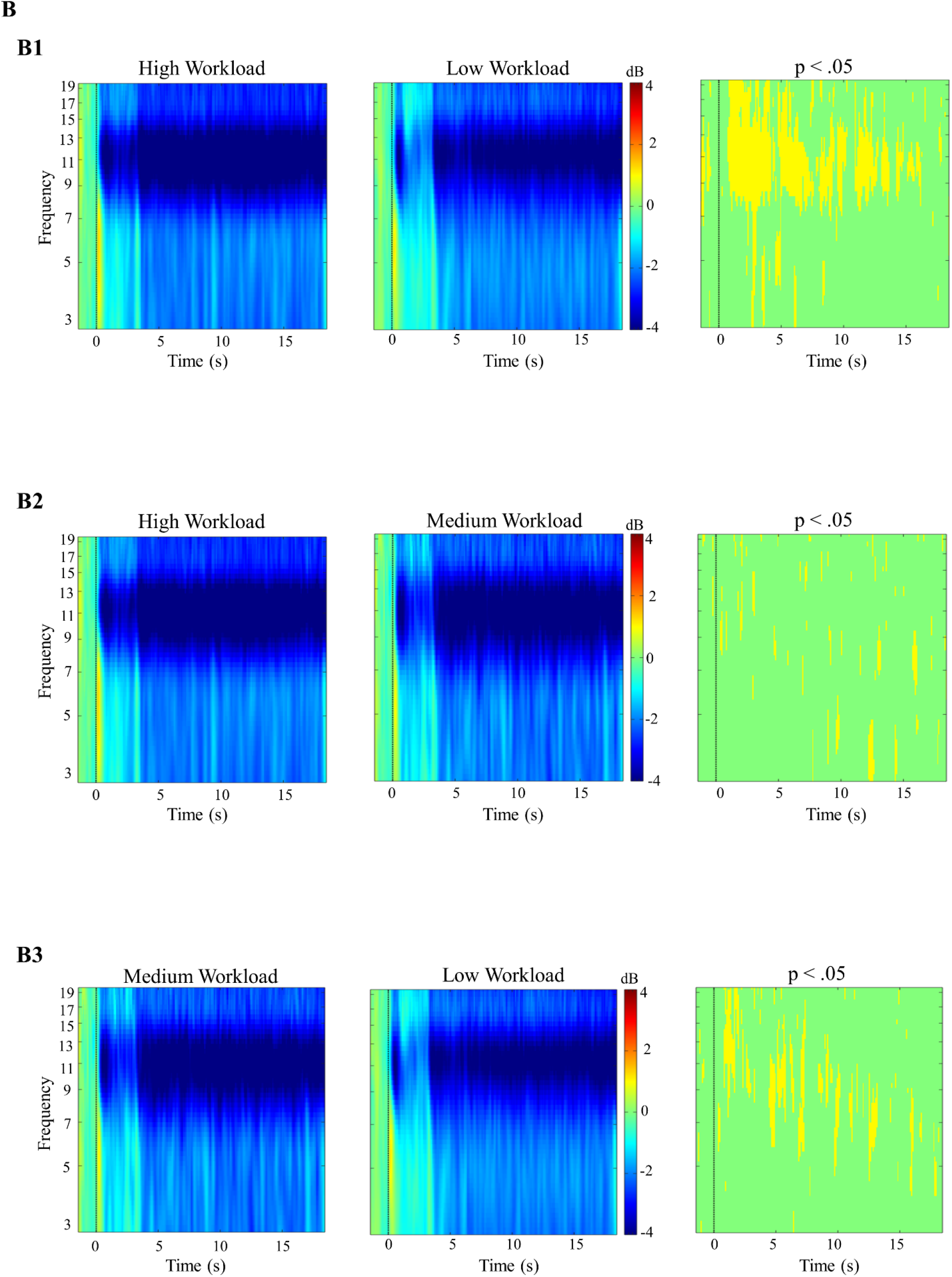
ERSP changes during the tracking task at the (A) Frontal and (B) Occipital Cluster. (A1) shows the ERSP changes at the frontal cluster during high (first panel) and low (second panel) workload conditions and the third panel shows the statistically significant difference between high and low workload conditions (p < .05). (A2) shows the ERSP changes at the frontal cluster during high (first panel) and medium (second panel) workload conditions and the third panel shows the statistically significant difference between high and medium workload conditions (p < .05). (A3) shows the ERSP changes at the frontal cluster during medium (first panel) and low (second panel) workload conditions and the third panel shows the statistically significant difference between medium and low workload conditions (p < .05). (B1) shows the ERSP changes at the occipital cluster during high (first panel) and low (second panel) workload conditions and the third panel shows the statistically significant difference between high and low workload conditions (p < .05). (B2) shows the ERSP changes at the occipital cluster during high (first panel) and medium (second panel) workload conditions and the third panel shows the statistically significant difference between high and medium workload conditions (p < .05). (B3) shows the ERSP changes at the occipital cluster during medium (first panel) and low (second panel) workload conditions and the third panel shows the statistically significant difference between medium and low workload conditions (p < .05).

Figure 6 illustrates the frontal, parietal and occipital clusters’ ERSP changes for three workload conditions in the collision prediction task. Statistical analysis on ERSP changes of the frontal cluster showed a significant increase in theta power with increasing levels of workload (Figure 6(A)). The ERSP changes at the parietal cluster (Figure 6(B)) revealed a significant increase in the theta power and a significant decrease in the alpha power with increasing level of workload. The ERSP changes at the occipital cluster (Figure 6(C)) revealed a significant increase in the delta and theta power with increasing workload.

**Figure 6:**
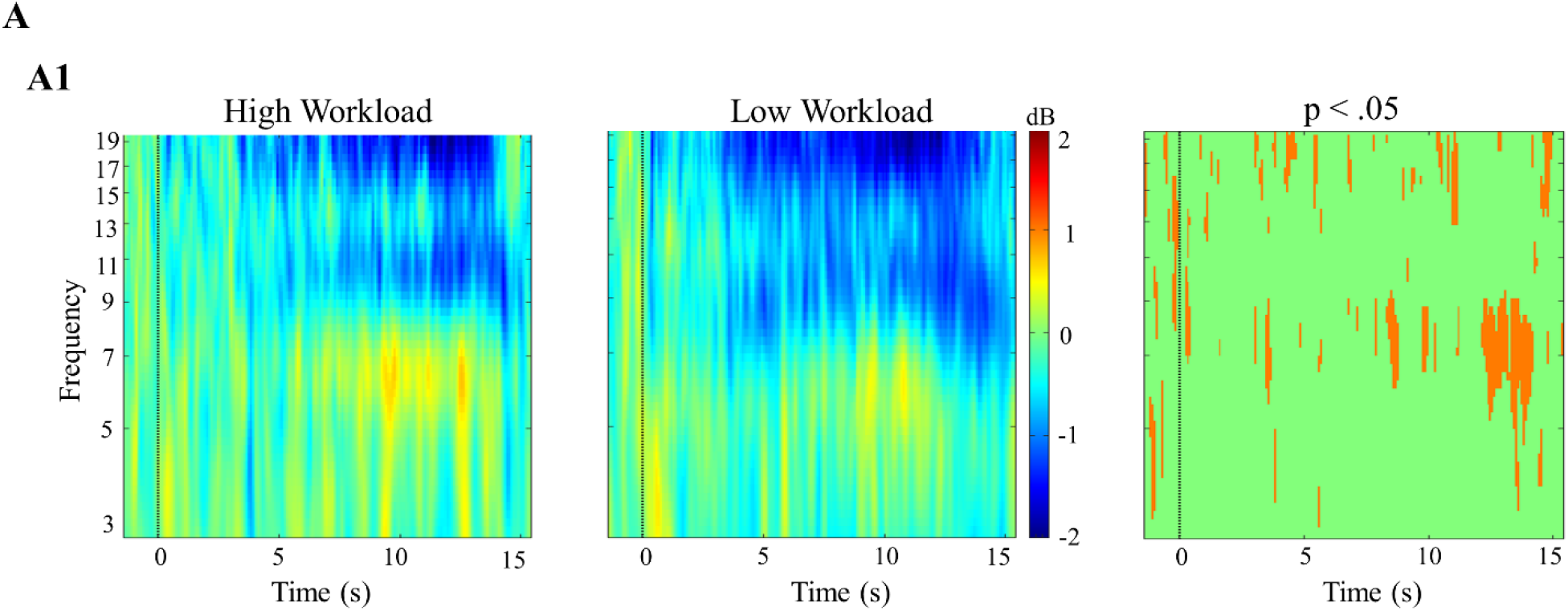
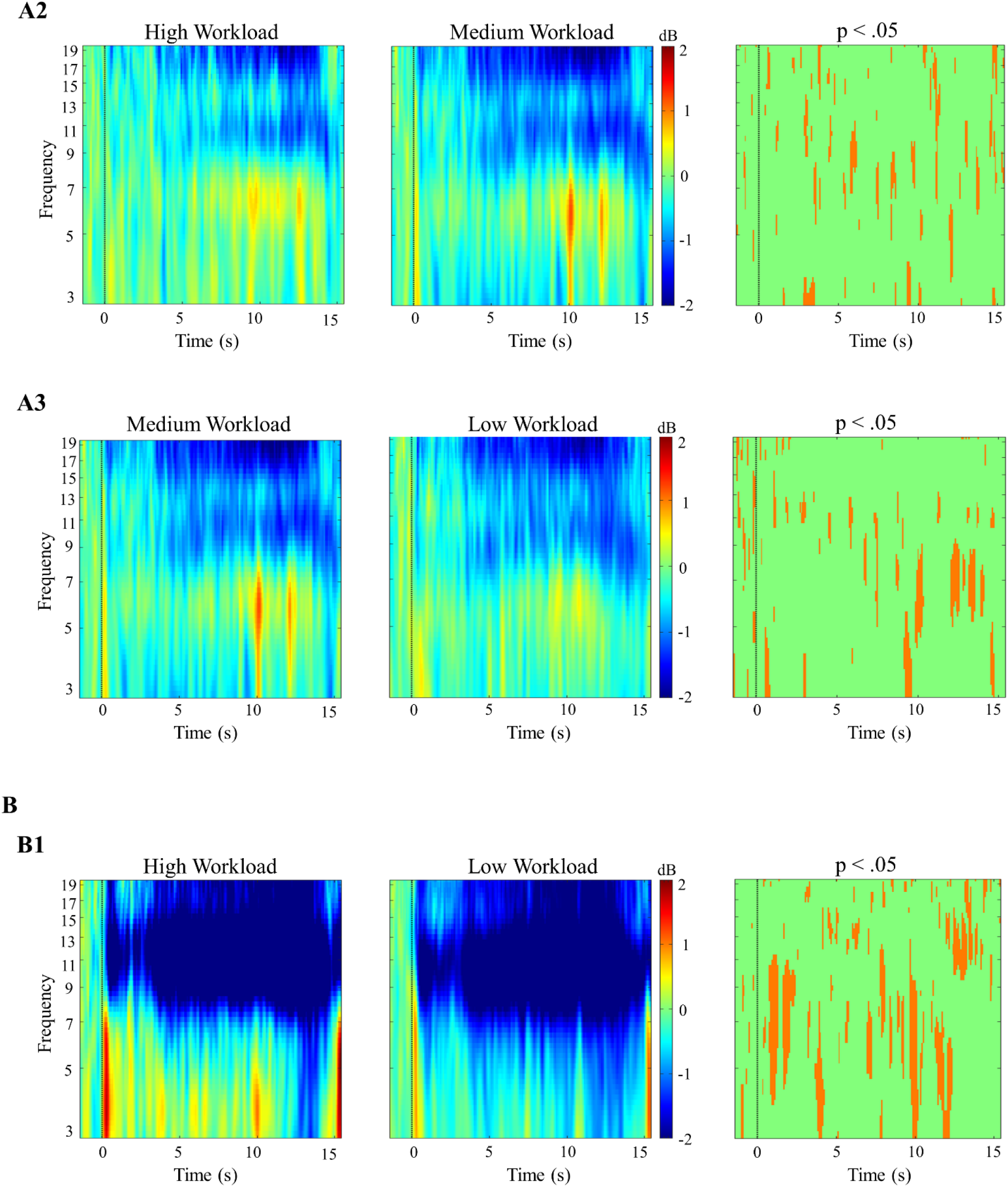
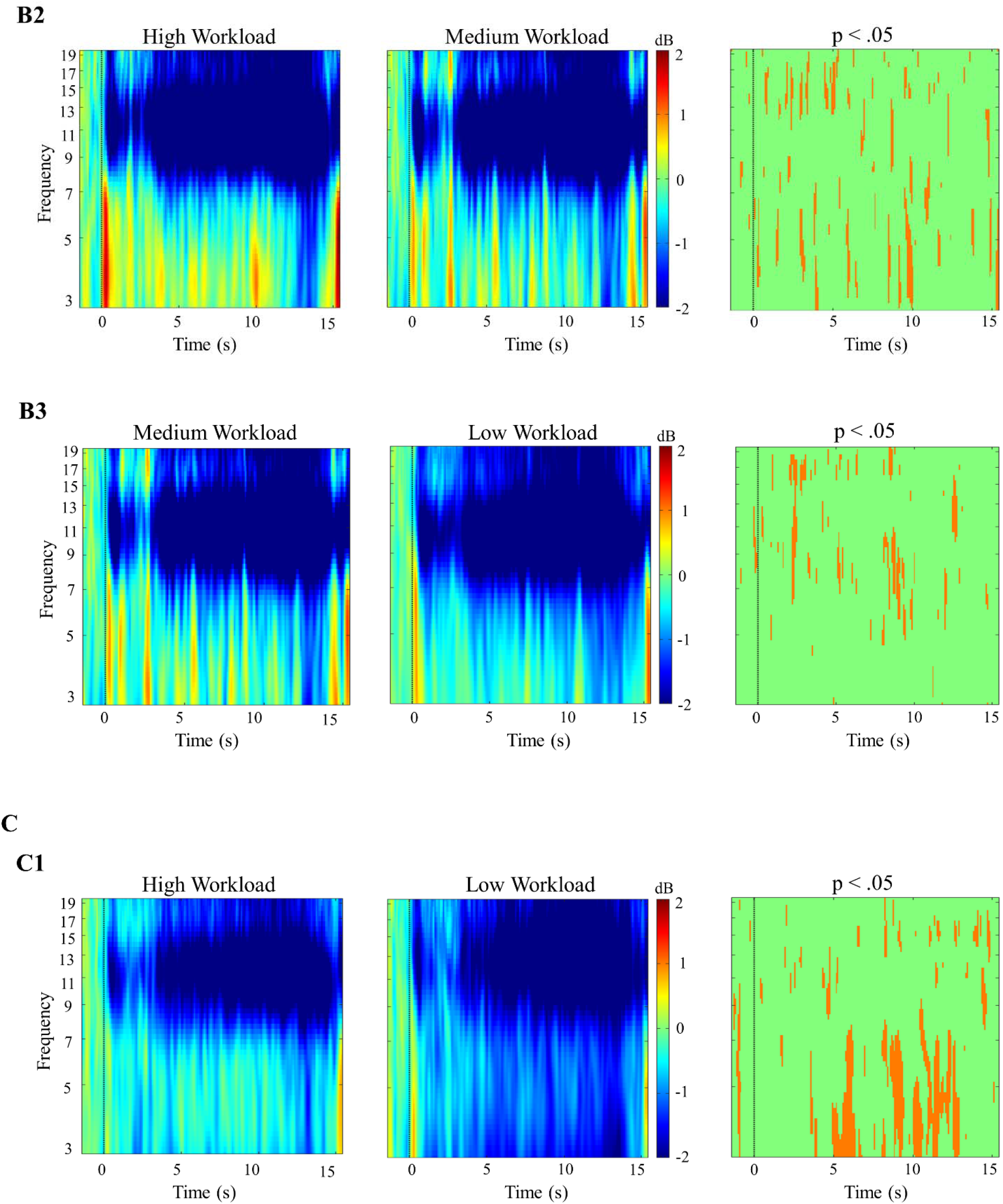
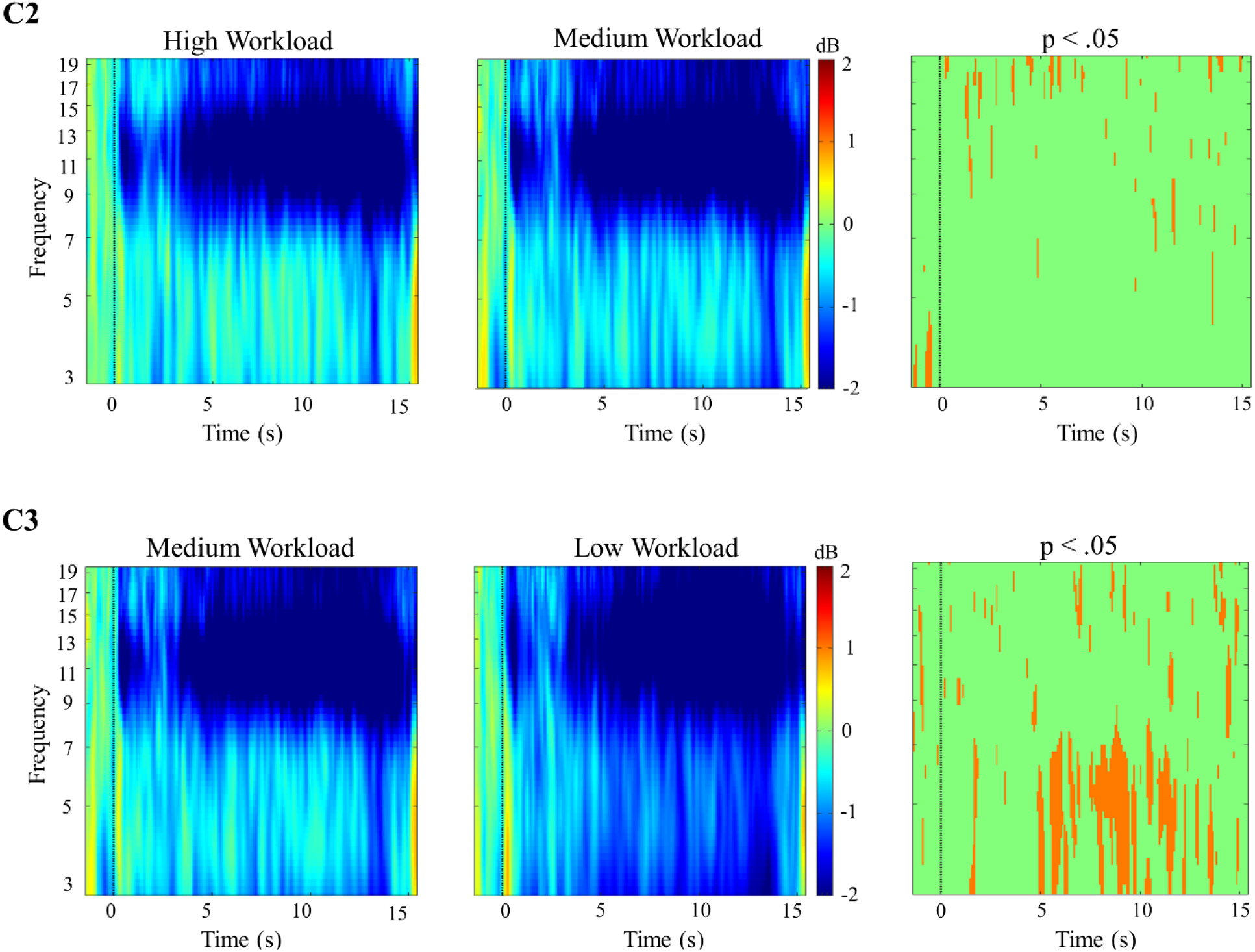
ERSP changes during the collision prediction task at the (A) Frontal, (B) Parietal, (C) Occipital Cluster. (A1) shows the ERSP changes at the frontal cluster during high (first panel) and low (second panel) workload conditions and the third panel shows the statistically significant difference between high and low workload conditions (p < .05). (A2) shows the ERSP changes at the frontal cluster during high (first panel) and medium (second panel) workload conditions and the third panel shows the statistically significant difference between high and medium workload conditions (p < .05). (A3) shows the ERSP changes at the frontal cluster during medium (first panel) and low (second panel) workload conditions and the third panel shows the statistically significant difference between medium and low workload conditions (p < .05). (B1) shows the ERSP changes at the parietal cluster during high (first panel) and low (second panel) workload conditions and the third panel shows the statistically significant difference between high and low workload conditions (p < .05). (B2) shows the ERSP changes at the parietal cluster during high (first panel) and medium (second panel) workload conditions and the third panel shows the statistically significant difference between high and medium workload conditions (p < .05). (B3) shows the ERSP changes at the parietal cluster during medium (first panel) and low (second panel) workload conditions and the third panel shows the statistically significant difference between medium and low workload conditions (p < .05). (C1) shows the ERSP changes at the occipital cluster during high (first panel) and low (second panel) workload conditions and the third panel shows the statistically significant difference between high and low workload conditions (p < .05). (C2) shows the ERSP changes at the occipital cluster during high (first panel) and medium (second panel) workload conditions and the third panel shows the statistically significant difference between high and medium workload conditions (p < .05). (C3) shows the ERSP changes at the occipital cluster during medium (first panel) and low (second panel) workload conditions and the third panel shows the statistically significant difference between medium and low workload conditions (p < .05).

#### Power Spectral Density Changes with Mental Workload

Figure 7(A1) illustrates that frontal theta PSD increased significantly with increasing levels of workload in the tracking task [F(2, 46) = 50.931, p < .001, η_p_^2^ = .822]. As shown in Figure 7(A2), the results of one-way repeated-measures ANOVA showed that occipital alpha PSD decreased significantly with increasing workload of the tracking task [F(2, 46) = 24.780, p < .001, η_p_^2^ = .693].

**Figure 7:**
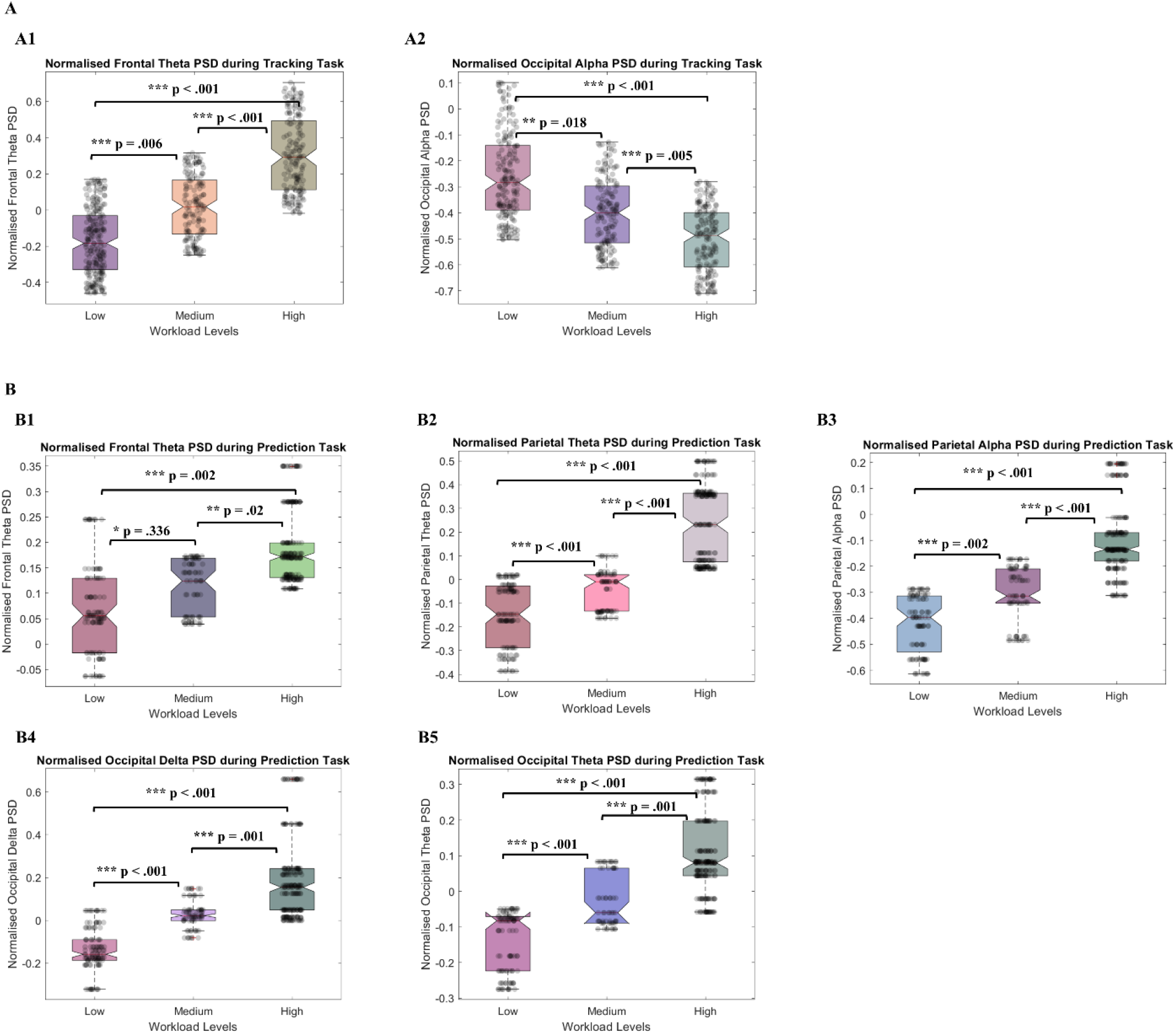
(A) Normalized Power Spectral Density at the Frontal and Occipital ICs selected in the Frontal and Occipital clusters for the tracking task. (A1) shows the normalised frontal theta PSD in the low, medium, and high workload conditions. (A2) shows the normalised occipital alpha PSD for low, medium, and high workload condition for the tracking task. (B) shows the normalized Power Spectral Density at the Frontal, Parietal and Occipital ICs selected in the Frontal, Parietal and Occipital cluster for the collision prediction task. (B1) shows the mean frontal theta PSD in the low, medium, and high workload conditions. (B2) shows the mean parietal theta PSD for the three levels of workload. (B3) shows the mean parietal alpha power for different workload conditions and (B4) shows the mean occipital delta PSD in the low, medium, and high workload conditions. (B5) shows the mean occipital theta PSD for the three levels of workload condition in collision prediction task.

For the collision prediction task, the frontal cluster’s ICs showed significant increase in theta PSD with increasing workload (Figure 7B(1)) according to the one-way repeated-measures ANOVA [F(2, 46) = 8.570, p = .001, η_p_^2^ = .271]. However, the parietal cluster’s IC’s spectral power showed a significant increase in the theta frequency band [F(2, 46) = 47.764, p < .001, η_p_^2^ = .675] and a significant decrease in the alpha band [F(2, 46) = 38.639, p < .001, η_p_^2^ = .627] with increasing workload, as shown in Figure 7(B2) and Figure 7(B3). One-way repeated-measures ANOVA results showed that occipital delta [F(1.563, 35.951) = 35.321, p < .001, η_p_^2^ = .606] and theta [F(2, 46) = 39.101, p < .001, η_p_^2^ = .630] power increased significantly with increasing workload in the collision prediction task, as shown in Figure 7(B4) and 7(B5).

#### Eye activity changes with mental workload

As shown in Figure 8(A), pupil size increased with the increasing workload for both tracking [F(2, 38) = 13.205, p < .001,η_p_^2^ = .410] and collision prediction tasks [F(2, 46) = 9.276, p < .001, η_p_^2^= .287]. The number of blinks during tracking and collision prediction tasks decreased with the increasing workload, as shown in Figure 8(B). One-way repeated-measure ANOVA was conducted to study the effect of workload variations on the number of blinks, which revealed significant variations in the number of blinks during the tracking task for different workload conditions [F(2, 46) = 3.624, p = .035, η_p_^2^ = .136]. The effect of workload on the number of blinks in the collision prediction task was analysed using one-way repeated-measure ANOVA. It showed a significant variation in the number of blinks [F(2, 46) = 18.586, p < .001, η_p_^2^ = .447].

**Figure 8:**
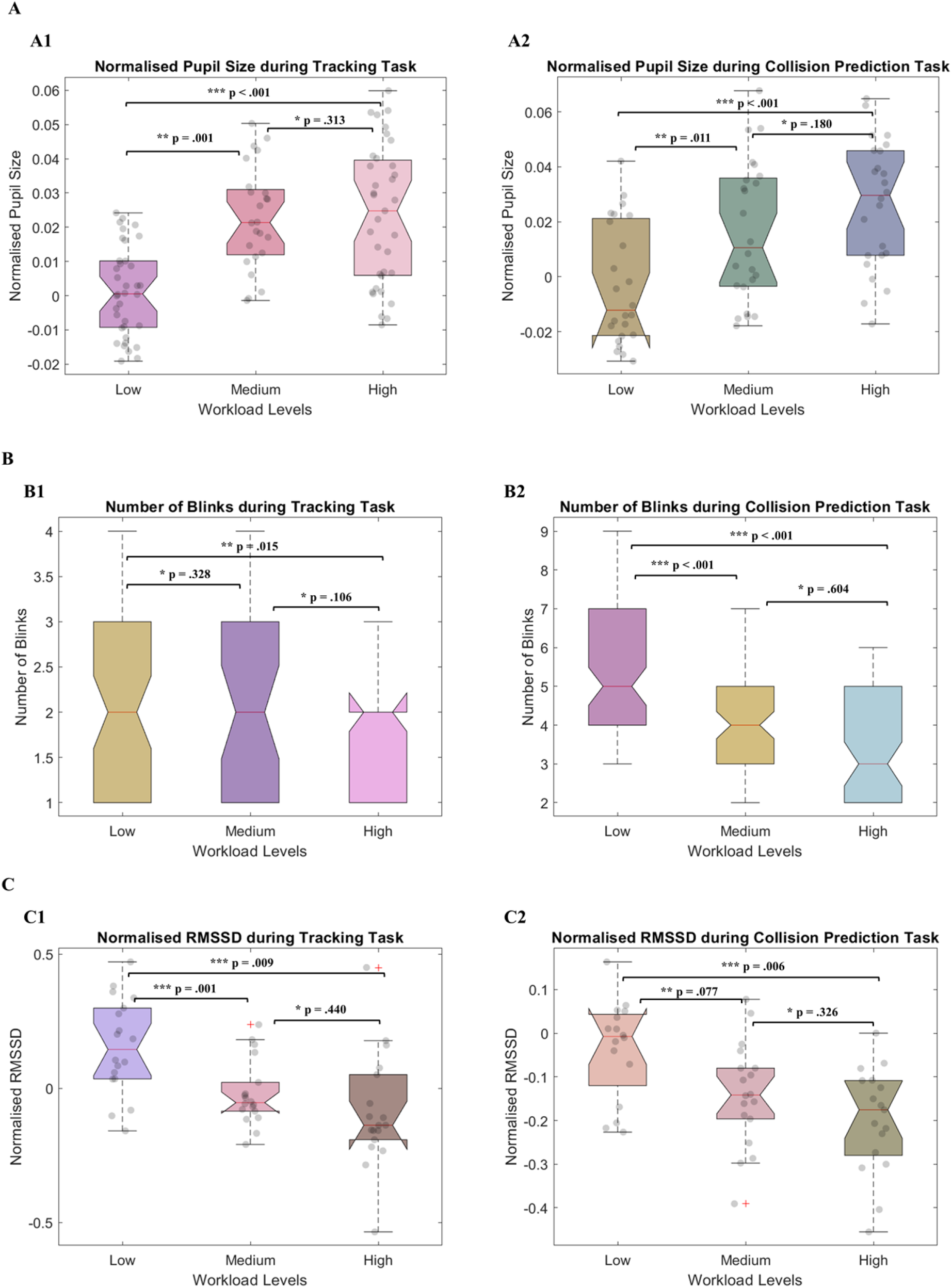
(A) shows the normalized pupil size of all the participants shows a positive trend with the increasing workload. (A1) Normalised pupil size in the three workload conditions of the tracking task. (A2) Normalised pupil size during low, medium, and high workload conditions for the collision prediction task. (B) shows the negative trend in the number of blinks with the increasing workload. (B1) Number of blinks during different workload conditions of the tracking task. (B2) Number of blinks during the collision prediction task decreases with increasing level of workload. (C) shows the declining trend in the normalized RMSSD of all the participants with the increasing workload. (C1) Normalised RMSSD all the participants in the low, medium, and high workload conditions of the tracking task. (C2) Normalised RMSSD during collision prediction task for the three levels of workload.

#### Heart Rate Variability (RMSSD) changes with Mental Workload

Figure 8(C) shows the RMSSD decreased significantly with increasing workload conditions of the tracking and collision prediction task. For the tracking task, there was a significant change in the RMSSD for the different workload conditions, as shown by the one-way repeated-measures ANOVA [F(2, 34) = 10.171, p < .001, η_p_^2^ = .374]. Results from one-way repeated-measures ANOVA shows that in the collision prediction task, there was a significant change in the RMSSD for different workload conditions [F(2, 44) = 4.279, p = .022, η_p_^2^ = .201].

#### Multiple Regression Results

Multiple regression was carried out to investigate whether EEG, eye activity and HRV metrics of workload could significantly predict the performance in the tracking task. The results of the regression indicated that the model explained 54.3% of the variance and that the model was a significant predictor of the tracking performance, F(3, 67) = 26.543, p < .001. While EEG metrics (p = .001) and eye activity (p < .001) contributed significantly to the model, HRV metrics did not (p = .125). The final predictive model was:

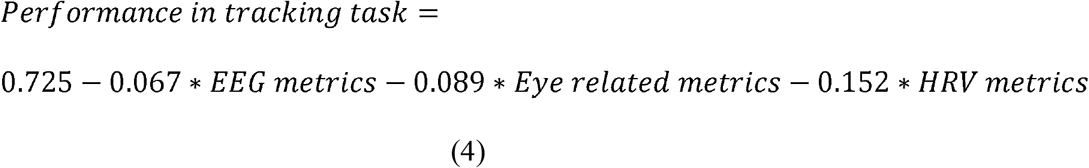

In order to determine whether EEG, eye activity and HRV metrics could significantly predict the performance in collision prediction task, we conducted multiple regression analysis. The results of the regression indicated that the model explained 61.7% of the variance and that the model was a significant predictor of the performance in the collision prediction task, F(3, 68) = 24.324, p < .001. While eye activity (p = .02) and EEG metrics (p < .001) contributed significantly to the model, HRV metrics did not (p = .443). The final predictive model was:

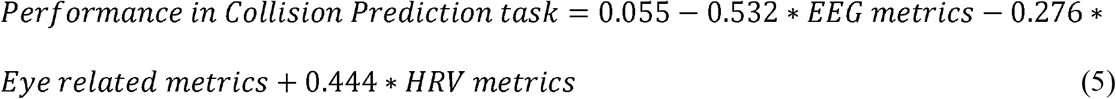

## Discussion

In this study, we designed two simplified tasks based on ATC: tracking and collision prediction tasks. Although both these tasks represent the basic tasks that ATC operators routinely perform, we considered them separately to untangle the differences in the physiological response to workload variations in these tasks.

In order to study workload effects of increasing air traffic, the mental workload in both these tasks was manipulated by varying the number of dots. It was observed that the performance in the tracking and collision prediction task deteriorated significantly with increasing levels of workload. Hence, we can confirm that the workload manipulation (by varying the number of dots) in both tracking and collision prediction tasks was successful in eliciting significant performance variations (H1).

In order to assess the mental workload, EEG, eye activity and BVP data were recorded while the participants performed the tasks. The tracking task demands allocation of attentional resources to keep track of one, three or five tracking dots moving randomly among distractor dots. Working memory load is sensitive to increased allocation of attentional resources and is reflected by increases in frontal theta power (Klimesch et al., 1998; Klimesch, 1999; Gevins and Smith, 2000). In the tracking task, we observed an increase in the frontal theta power, which confirms that increased working memory load was experienced with increasing workload levels. Tracking dots moving among distractor dots also entails working memory mechanisms related to relevant item maintenance and increases in the memory load. This working memory mechanism was reflected by a decrease in the alpha power (Gevins et al., 1997; Wilson, 2002 and Puma et al., 2018). The alpha power is also known to decrease with increased memory load (Fournier et al., 1999; Smith et al., 2001; Ryu and Myung, 2005) and task difficulty (Sterman and Mann, 1995; Ota et al., 1996). Our findings also substantiate this working memory mechanism as the occipital alpha power decreases with increasing workload levels in the tracking task.

In the collision prediction task, anticipating the trajectory of the dots and predicting whether the dots would collide requires attention and internal concentration. Delta power is an indicator of attention or internal concentration in mental tasks, and it has been reported to increase with the increase in workload (Sterman and Mann, 1995; Harmony et al., 1996; Wilson, 2002). Our results demonstrate an increase in the delta power at the occipital sites, which validates that there is an increased allocation of attentional resources with increasing levels of workload in the collision prediction task. Additionally, keeping a tab on the trajectory of six, 12 or 18 eight dots adds to the memory load in the participants. Several studies have shown that theta power is correlated with memory load (Jensen and Tesche, 2002; Jacobs et al., 2006) and working memory capacity (Klimesch, 1996; Klimesch, 1999; Sauseng et al., 2010). In collision prediction task, our results reveal a significant increase in the theta power at the frontal, parietal and occipital clusters, confirming an increase in memory load with increasing levels of workload. Furthermore, our results indicate that with increasing levels of workload, there is a decrease in parietal alpha power. This observed alpha band desynchronisation with the increasing workload is related to relevant item maintenance in the working memory (Sterman and Mann, 1995; Gevins et al., 1997; Wilson, 2002; Puma et al., 2018) and is known to decrease with increased memory load (Fournier et al., 1999; Smith et al., 2001; Ryu and Myung, 2005) and task difficulty (Sterman and Mann, 1995; Ota et al., 1996). However, in the collision prediction task, the most significant decrease in the parietal alpha power was observed a few seconds before the collision. It might be related to the increase in the experienced time pressure (Slobounov et al., 2000) as the participants attempt to identify and click on the colliding pair of dots before the collision happens.

We also explored eye-related metrics and HRV metrics during workload variations. Eye activity data was transformed to pupil size and blink rate. Pupil size increased significantly with the increasing workload in both tracking and collision prediction tasks. The number of blinks also reduced considerably with the increasing workload in both tasks. Pupil size is a reliable measure of workload (Marquart et al., 2015) as it dilates with increasing workload. Recarte et al., 2008 show that blink inhibition happens in higher workload conditions and so, the blink rate is inversely correlated with the attentional levels and workload experienced by the operator (Brookings et al., 1996, Wilson, 2002, Widyanti et al., 2017). RMSSD was found to be negatively correlated with the mental workload in both tasks. This decrease in RMSSD with the increasing workload is widely reported in the literature (Mehler et al., 2011, Heine et al., 2017).

Our results show that EEG power spectra at the frontal, parietal and occipital areas, eye activity and HRV metrics can reliably and accurately assess the mental workload of the participants in both tasks. Hence, our second hypothesis (H2) is proved to be true for both tracking and collision prediction tasks. Relating to our third hypothesis (H3), the multiple regression results showed that the performance in the tracking and collision prediction tasks could be predicted from the EEG, eye related and HRV metrics.

Our results also indicate that even though eye activity and HRV metrics are sensitive to task load variations, they may not provide any valuable information on the task that causes the variations in workload. However, the EEG measures were found to be not just sensitive to the workload variations but also the task type. The increases in workload in the tracking task was reflected by the increase in frontal theta power and decrease in occipital alpha power. No significant changes were observed in the parietal theta, alpha, occipital delta, or theta power with the increasing workload in the tracking task. In the collision prediction task, the increase in workload was correlated with the increases in frontal theta, parietal theta, occipital delta and theta power and a decrease in parietal alpha power. No significant variation was observed in the occipital alpha power during the collision prediction task. The neurometrics correlated with the variations in the workload of tracking and collision prediction tasks are different, which proves that our fourth hypothesis (H4) is true. Therefore, neurometrics can help identify the task contributing to the increase in workload in complex ATC environments at a time instant and define the strategies that can be used by the workload adaptive system to mitigate this increase. These results provide evidence that the use of EEG measures in a closed-loop adaptive system can not only aid the decision of “when” but also “what” form of automation to deploy to mitigate the workload variations in operators. Hence, the results presented here contribute to the development of adaptive strategies essential for the design of intelligent closed-loop mental workload adaptive ATC systems.

## Conclusion

In order to elucidate the impact of basic task load variations that comprise the load variations in complex ATC tasks, we separately designed two basic ATC tasks: tracking and collision prediction tasks. EEG spectral power, eye and HRV correlates to mental workload variations for tracking and collision prediction tasks of air traffic controllers are successfully unravelled. The differences in neural response to increased workload in the tracking and collision prediction task indicate that these neural measures are sensitive to variations and type of mental workload and their potential utility in not just deciding “when” but also “what” to adapt, aiding the development of intelligent closed-loop mental workload aware systems. This investigation of physiological indices of workload variation in the basic ATC tasks has applicability to the design of future adaptive systems that integrate neurometrics in deciding the form of automation to be used to mitigate the variations in workload in complex ATC systems.

## Acknowledgements

This work was supported in part by the Australian Research Council (ARC) under Discovery Grant DP180100670 and Discovery Grant DP180100656; in part by the Australia Defence Innovation Hub under Contract P18-650825; U.S. Office of Naval Research Global through Cooperative Agreement under Grant ONRG-NICOP-N62909-19-1-2058; and in part by the NSW Defence Innovation Network and NSW State Government of Australia under Grant DINPP2019 S1-03/09.

## Key points

- Workload variation in tracking and collision prediction tasks was reliably assessed using EEG, eye activity and HRV metrics.
- The performance in tracking and collision prediction tasks can be predicted based on the measured physiological signals.
- Neurometrics of the workload variations in the tracking and collision prediction tasks are distinct across tasks.

## Supplementary Material

**Figure 1.**
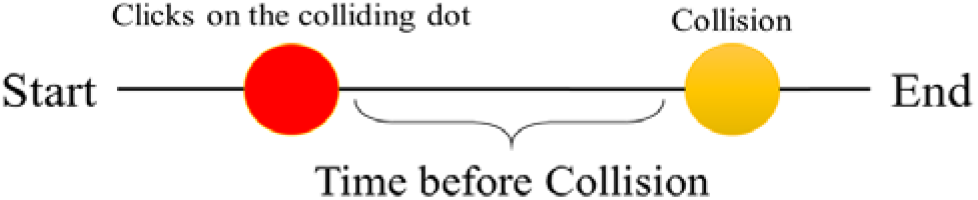
A schematic diagram describing how time before collision was calculated in the collision prediction task

**Figure 2.**
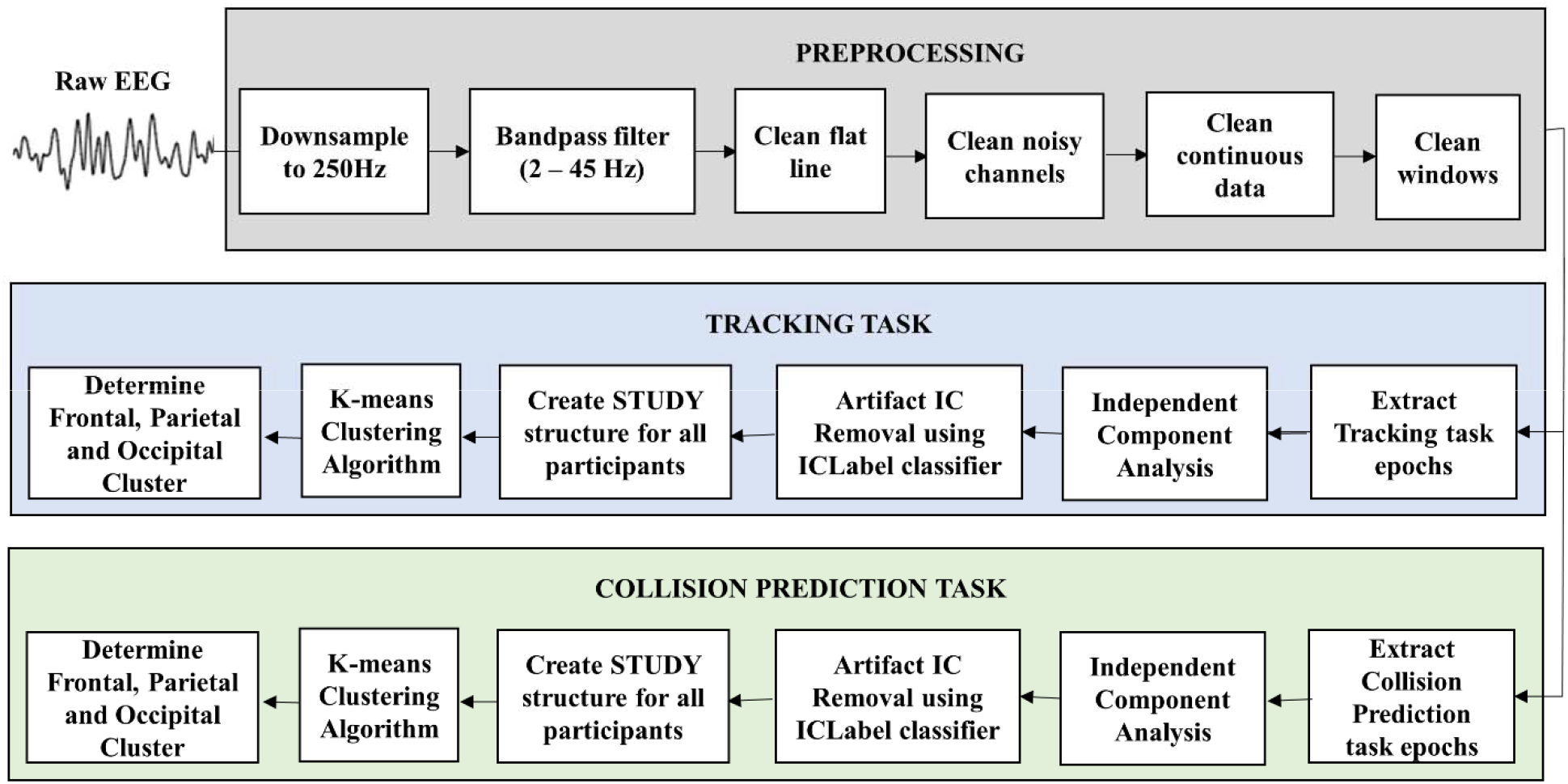
The EEG preprocessing and processing pipeline used for tracking and collision prediction tasks.

